# Comparative analysis of myoglobin in Cetaceans and humans reveals novel regulatory elements and evolutionary flexibility

**DOI:** 10.1101/2023.04.10.536305

**Authors:** Charles Sackerson, Vivian Garcia, Nicole Medina, Jessica E. Maldonado, John Daly, Rachel Cartwright

## Abstract

Cetacea and other diving mammals have undergone numerous adaptations to their aquatic environment, among them high levels of the oxygen-carrying intracellular hemoprotein myoglobin in skeletal muscles, especially those responsible for swimming. Hypotheses regarding the mechanisms leading to these high myoglobin levels often invoke exercise, hypoxia, and other physiological pathways that impact gene regulation, and increase myoglobin gene expression. Here we explore an alternative hypothesis: that Cetacean myoglobin genes have evolved high levels of transcription driven by the intrinsic developmental mechanisms that drive muscle cell differentiation. We have used luciferase assays in differentiated C2C12 cells to test this hypothesis. Contrary to our hypothesis, we find that the myoglobin gene from the minke whale, *Balaenoptera acutorostrata*, has a low level of expression, less than 10% that of humans. This low expression level is broadly shared among Cetaceans, and Artiodactylans. Previous work on regulation of the human gene has identified a core muscle-specific enhancer comprised of two regions, the “AT element” and a C-rich sequence 5’ of the AT element termed the “CCAC-box”. Comparative analysis of the minke whale gene supports the importance of the AT element, but the minke whale CCAC-box ortholog has little effect. Instead, critical positive input has been identified in a G-rich region 3’ of the AT element. Also, a conserved E-box in exon 1 positively affects expression, despite having been assigned a repressive role in the human gene. Last, a novel region 5’ of the core enhancer has been identified, which we hypothesize may function as a boundary element. These results illustrate regulatory flexibility during evolution. They also support the hypothesis that the induction of transcription by physiological signaling pathways, and evolved protein stability are important in leading to the high myoglobin protein levels found in Cetaceans.

## Introduction

The Cetacea diverged from their terrestrial relatives about 50 million years ago [1–3], and have since undergone numerous adaptations to their aquatic lifestyle. Among these is high levels of the intracellular hemoprotein myoglobin (MB) in skeletal muscle [4], especially in those muscles required for swimming. Myoglobin stores oxygen and assists in its facilitated diffusion to supply oxygen in muscle tissue during extended dives [5]. Myoglobin also manages reactive oxygen and nitrogen species [6], regulates mitochondrial respiration [7], and carries out other functions relevant to a relatively anoxic muscle environment [8]. These functions all contribute to muscle fitness and functionality during exercise in anoxic /oxygen limited environments.

We are interested in the mechanisms underlying the evolution of high levels of myoglobin in Cetacean muscles. Traditionally, evolution has focused on changes in amino acid sequence, but recent studies have highlighted the importance of changes in gene regulation as drivers of evolutionary change [9–11]. Therefore, we hypothesized that the high myoglobin protein levels may be the result of evolutionary adaptations affecting the regulation of the myoglobin gene, which in turn lead to high rates of transcription. That is, even in the absence of induction by exercise or other physiological influences, the constitutive, or basal, level of transcription would be sufficient to lead to high myoglobin protein levels. This may be particularly relevant to myoglobin levels in neonates, which have not yet been influenced by exercise [12].

Whereas the regulatory sequences that control muscle-specific expression of the myoglobin gene have been well studied in humans and mice [13-16,reviewed in 17], little is known about the regulation of the Cetacean genes. A summary of the 5’ regulatory region of the human gene is shown in Fig 1. Of special note is a muscle-specific core enhancer that responds to the developmental signals that trigger muscle cell differentiation. The two components of this enhancer, the CCAC-box and the AT element, act synergistically to drive high levels of transcription [15,18]. Mutations in either reduce transcription to 10-20% of wild-type [14]. Based on these findings, we cloned the 5’ regulatory region from several species spanning the evolutionary range from Cetaceans to humans and assayed their transcriptional activity in the mouse myoblast cell line, C2C12, after differentiation to myotubes.

**Fig 1.**
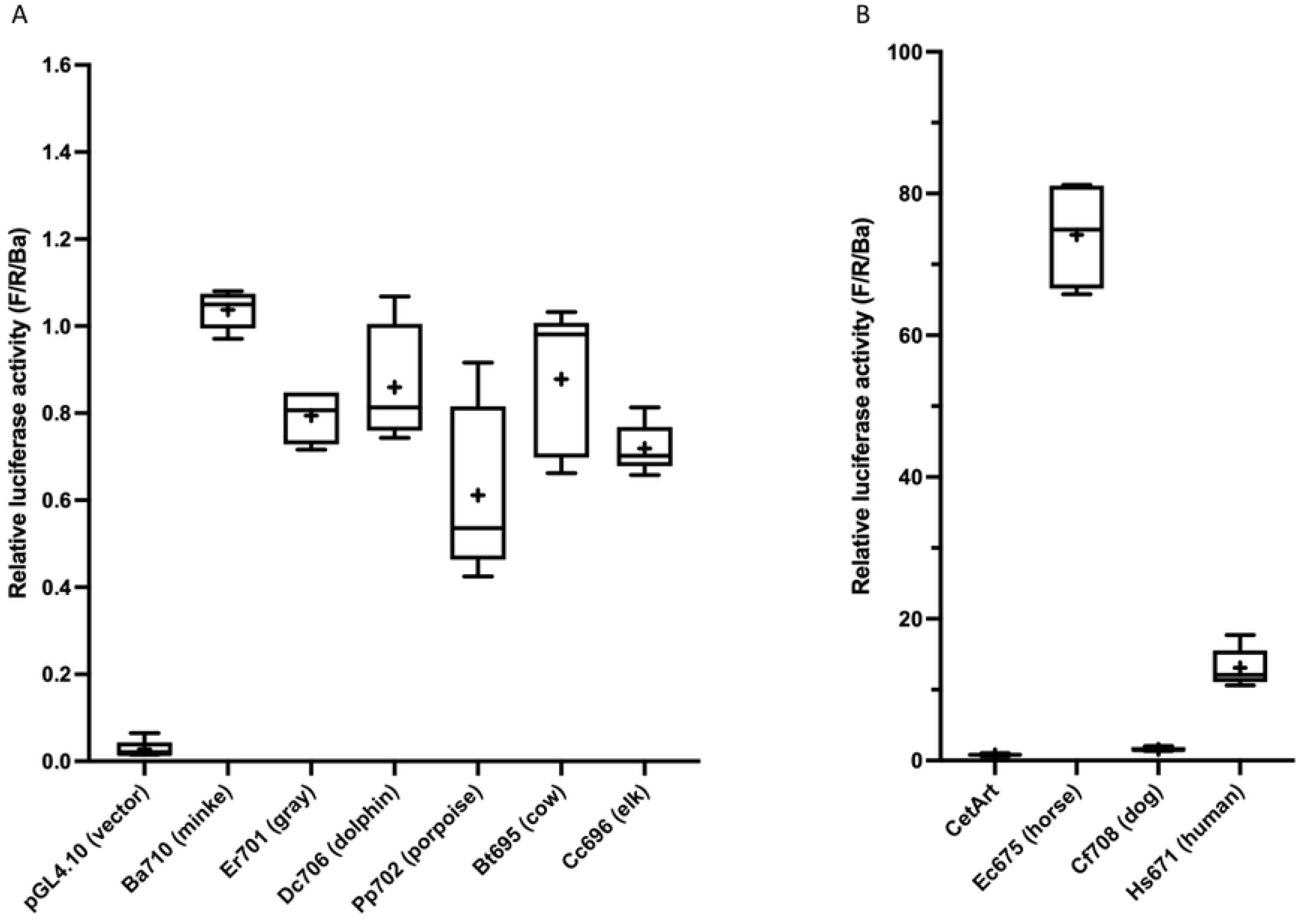
Summary of regulatory features in the myoglobin 5’ flanking region. DNA sequence elements identified previously or in this work are schematized, to scale. The top line represents the human (*Homo sapiens*, Hs) gene, the lower line the minke whale (*Balaenoptera acutorostrata*, Ba) gene. Hs Tss1 is the human major transcription start site (Tss) [19]; the Ba Tss is presumed to be at the same nucleotide. The arrows represent positive regulatory inputs unless otherwise noted. The “+” and “-” notation on the E-box3 arrows reflect its’ activating (positive) effect in the Ba gene and repressive (negative) effect in the Hs gene. The “X” on the Hs G-rich sequence arrow and the Ba CCAC-box arrow indicate a lack of effect. The “?” on the conserved Hs orthologue of the Ba449/412 region indicates that its effect on expression was not tested. DNaseI HS (p195602) indicates the extent of a DNaseI hypersensitive site identified in muscle cells by the ENCODE project as displayed in the UCSC Genome Browser [20,21] (see Fig 7 for further details).

To further understand the transcription of the Cetacean myoglobin gene, we then carried out a detailed dissection of the 5’ flanking region of the minke whale gene, as a model for Cetaceans in general. These results are summarized in Fig 1.

We report here two major findings from these studies. First, contrary to our hypothesis, we find that the Cetacean myoglobin genes do not have high transcriptional activity compared to the human gene, with expression levels at only about 1/16 on average, compared to that of humans. This implies that additional levels of regulation may be important such as the induction of increased transcription in response to physiological signaling pathways [12,22,23], and post-transcriptional mechanisms such as increased protein stability [24–26]. Second, considerable regulatory evolution has occurred in the myoglobin gene since a common ancestor with humans. Some regulatory elements such as the AT element are conserved in function, others such as the CCAC box have lost function, and novel regulatory elements are found that contribute to transcriptional activity. Cumulatively, these studies demonstrate mechanisms through which regulatory evolution occurs while conserving an essential regulatory strategy, in this case, muscle-specific gene expression.

## RESULTS

### Species survey

Previous studies of human (*Homo sapiens*, Hs) myoglobin (MB) regulation have indicated that the important regulatory inputs are present in about 700 nucleotides (nt) of 5’ flanking sequence [17,27]. We cloned and sequenced about 700 nt of 5’ flanking sequence from the MB genes of the baleen whales, minke whale, *B. acutorostrata* (Ba710, 710 nt of DNA 5’ of the translational start codon (ATG) from Ba), and gray whale, *Eschrichtius robustus* (Er701). Alignments with the published human sequence showed that these whale regions corresponded to 671 nt of Hs sequence; we therefore also cloned 671 nt of human sequence (Hs671) for comparative studies. These promoter regions were placed 5’ of the luciferase reporter in pGL4.10[luc2], transfected into the mouse myoblast cell line C2C12, and luciferase activity was measured after 4-6 days of differentiation to myotubes (see S1A File, Materials and Methods for details). We find that the whale promoters have relatively low activity compared to the human promoter, with the human promoter being >12-fold more active than Ba710 (Table 1, S2 File, Fig 2).

**Fig 2.**
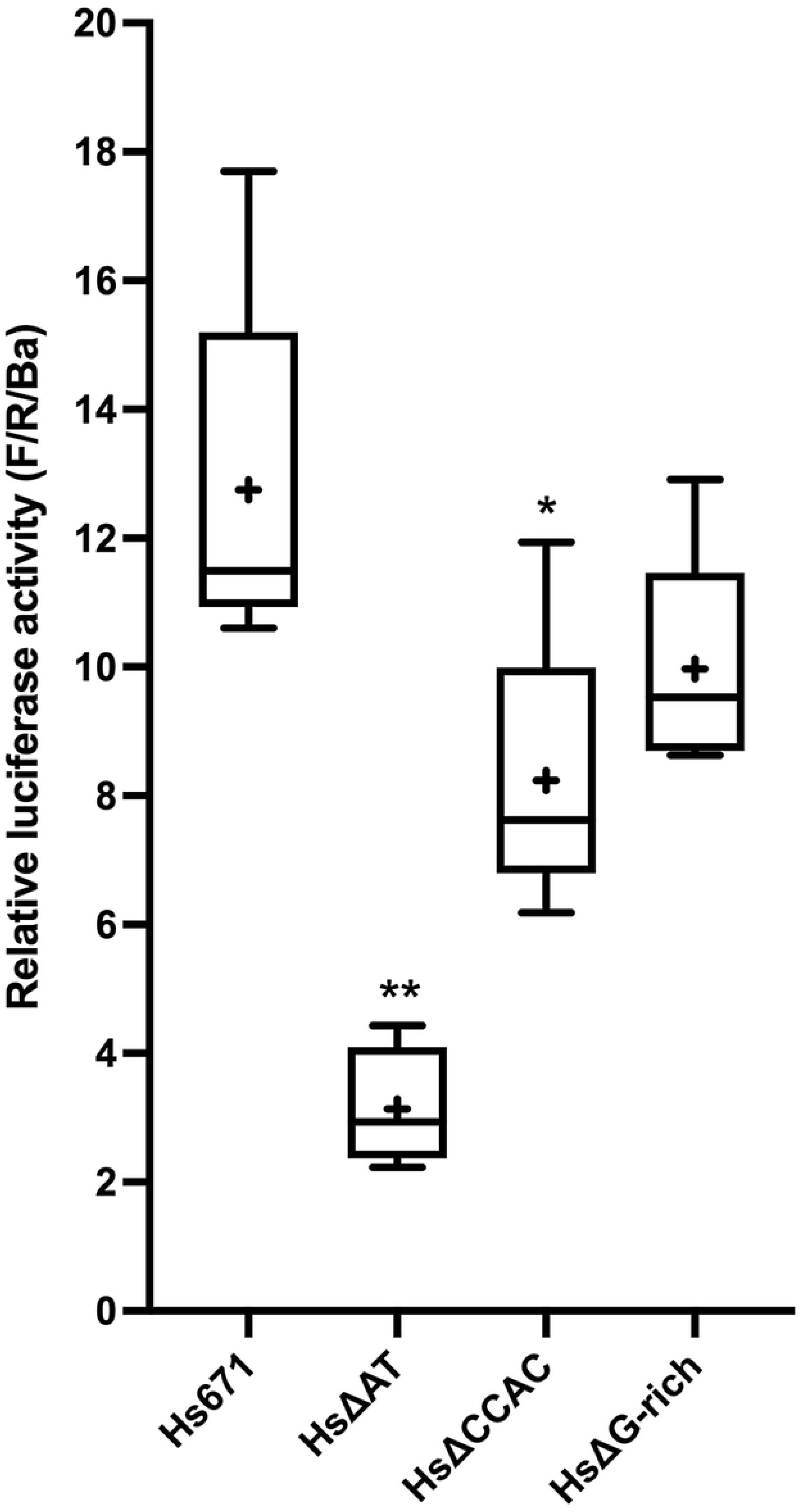
Species scan of MB promoter activity. (A) Box-and-whisker plots of selected Cetacean and Artiodactylan species, normalized as described for Table 1. The Y-axis shows activity compared to the Ba710 control included in each transfected plate. The “+” in each box is the sample mean. Clone designations are as described for Table 1. (B) The mean of ten Cetacean and Artiodactylan species (“CetArt”, S2 File) is compared to horses (Ec675: *E. caballus*), dogs (Cf708: *C. familiaris*), and humans (Hs671: *H. sapiens*). Note that the axes differ in scale in A and B.

**Table 1.**
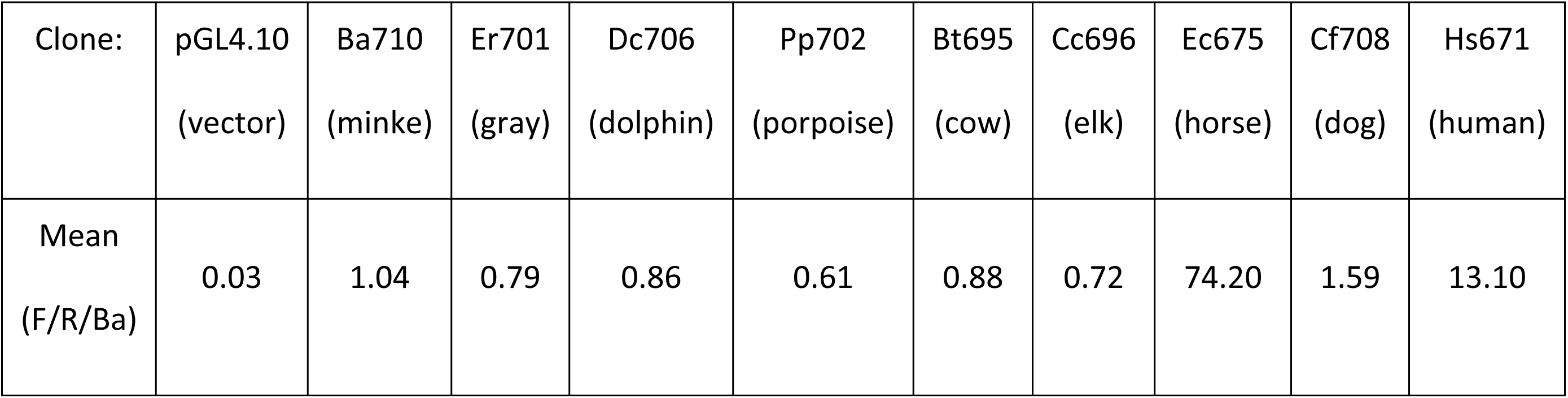
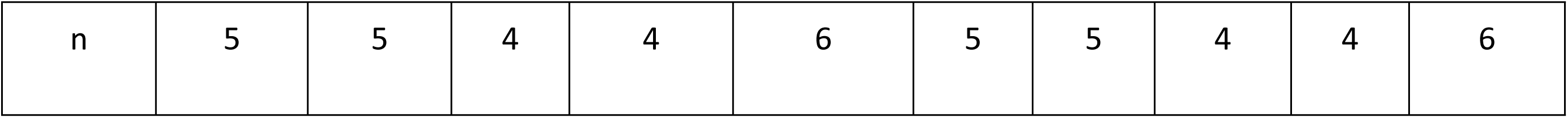
Species scan of MB promoter activity.

| Clone: | pGL4.10 | Ba710 | Er701 | Dc706 | Pp702 | Bt695 | Cc696 | Ec675 | Cf708 | Hs671 |
| --- | --- | --- | --- | --- | --- | --- | --- | --- | --- | --- |
|  | (vector) | (minke) | (gray) | (dolphin) | (porpoise) | (cow) | (elk) | (horse) | (dog) | (human) |
| Mean<br>(F/R/Ba) | 0.03 | 1.04 | 0.79 | 0.86 | 0.61 | 0.88 | 0.72 | 74.20 | 1.59 | 13.10 |
| n | 5 | 5 | 4 | 4 | 6 | 5 | 5 | 4 | 4 | 6 |

Mean activity of selected species, expressed as firefly luciferase (F) counts normalized first to a cotransfected renilla luciferase (R) internal standard, then secondly to a minke whale (Ba710) control included in duplicate in each transfection (F/R/Ba). The number of individual transfections, done in duplicate, is indicated as “n”; for example, n = 5 refers to 10 transfected wells in 5 independent experiments. pGL4.10 is empty vector. The Ba710 sample shown reflects independent transfections of Ba710, treated the same as the other samples. pGL4.10: empty vector, Ba710: *Balaenoptera acutorostrata*, Er701: *Eschrichtius robustus*, Dc706: *Delphinus capensis*, Pp702: *Phocoena phocoena*, Bt695: *Bos taurus*, Cc696: *Cervus canadensis*, Ec675: *Equus caballus*, Cf708: *Canis familiaris*, Hs671: *Homo sapiens*. Numbers indicate the size of each cloned promoter in base pairs, where the A of the translational initiating ATG = +1, and counting in the 5’ direction begins with the nucleotide immediately 5’ of the ATG, invariably a C in these species; thus, Ba710 has 710 nucleotides 5’ of the ATG. The transcription start site (Tss) is not used as +1 because various human Tss have been proposed in different sources, and the Tss from the other species have not always been experimentally determined.

We pursued these results by cloning and testing comparable regions from the toothed whales, common dolphin, *Delphinus capensis* (Dc706), and harbor porpoise, *Phocoena phocoen*a (Pp702); the closely-related terrestrial Artiodactylan (even-toed ungulate) species, cows, *Bos taurus* (Bt695), and elk, *Cervus canadensis* (Cc696); and more distantly-related species, horses, *Equus caballus* (Ec675, an odd-toed ungulate), and dogs, *Canis familiaris* (Cf708, a carnivore). Toothed whales, Artiodactylans, and dogs are not greatly different in activity from the Cetacea (Table 1, S2 File). The promoter from horses is highly active, >5-fold more so than humans, and >70-fold that of the minke whale (Table 1, S2 File). This species survey is summarized in Fig 2.

This survey demonstrates that the high levels of myoglobin protein in Cetacean muscle is unlikely to be due to a high constitutive level of transcriptional activity of the MB gene. We use the term “constitutive” to describe expression driven by differentiation of the C2C12 cells. Our assays do not address increases in expression “induced” by physiological signals such as calcium flux, hypoxia, or lipid availability.

### Exploration of the minke whale (*Ba*) myoglobin gene regulatory regions previously identified in the human (*Hs*) gene

The low activity of the Cetacean genes prompted us to explore in detail the transcriptional regulation of a model Cetacean MB gene, that of the minke whale, *B. acutorostrata* (Ba). Previous work on regulation of the Hs myoglobin gene identified a muscle-specific core enhancer composed of two sub-regions: an “AT element” and a “CCAC-box” [13–15,17]. In addition, an isolated E-box, “E-box3” [28] was identified in exon 1. The orthologous sequences from the Ba gene were identified by DNA sequence alignment and assessed for their contributions to gene regulation in differentiated C2C12 mouse myoblast cells. Binding sites for the transcription factor NFAT [29] are also able to be identified by sequence conservation or transcription factor searches, but our assays do not include manipulations that would activate the NFAT pathway, so these binding sites were not explicitly pursued.

#### AT element

The AT element consists of three separate transcription factor binding regions: an AT-rich region flanked on both sides by E-box sequences (Fig 3A). The AT-rich region has been shown to bind a critical activating transcription factor, MEF2 [18]. E-box sequences bind muscle-specific basic helix-loop-helix (bHLH) transcription factors such as MYOD and myogenin (MYOG) in cooperation with ubiquitous TCF3/E2A proteins such as E12 and E47 [30]. The AT-rich region is highly conserved across the species studied (S3A File). There are two nucleotide differences between Hs and Ba affecting the MEF2 binding consensus [31] (Hs: <u>CT</u>AAAATAG → Ba: <u>TC</u>AAAATAG), but the Ba AT element is still predicted (LASAGNA, rVISTA; see Materials and Methods) to bind MEF2. The two E-boxes are highly conserved in the species examined (except porpoises, see S3A File).

**Fig 3.**
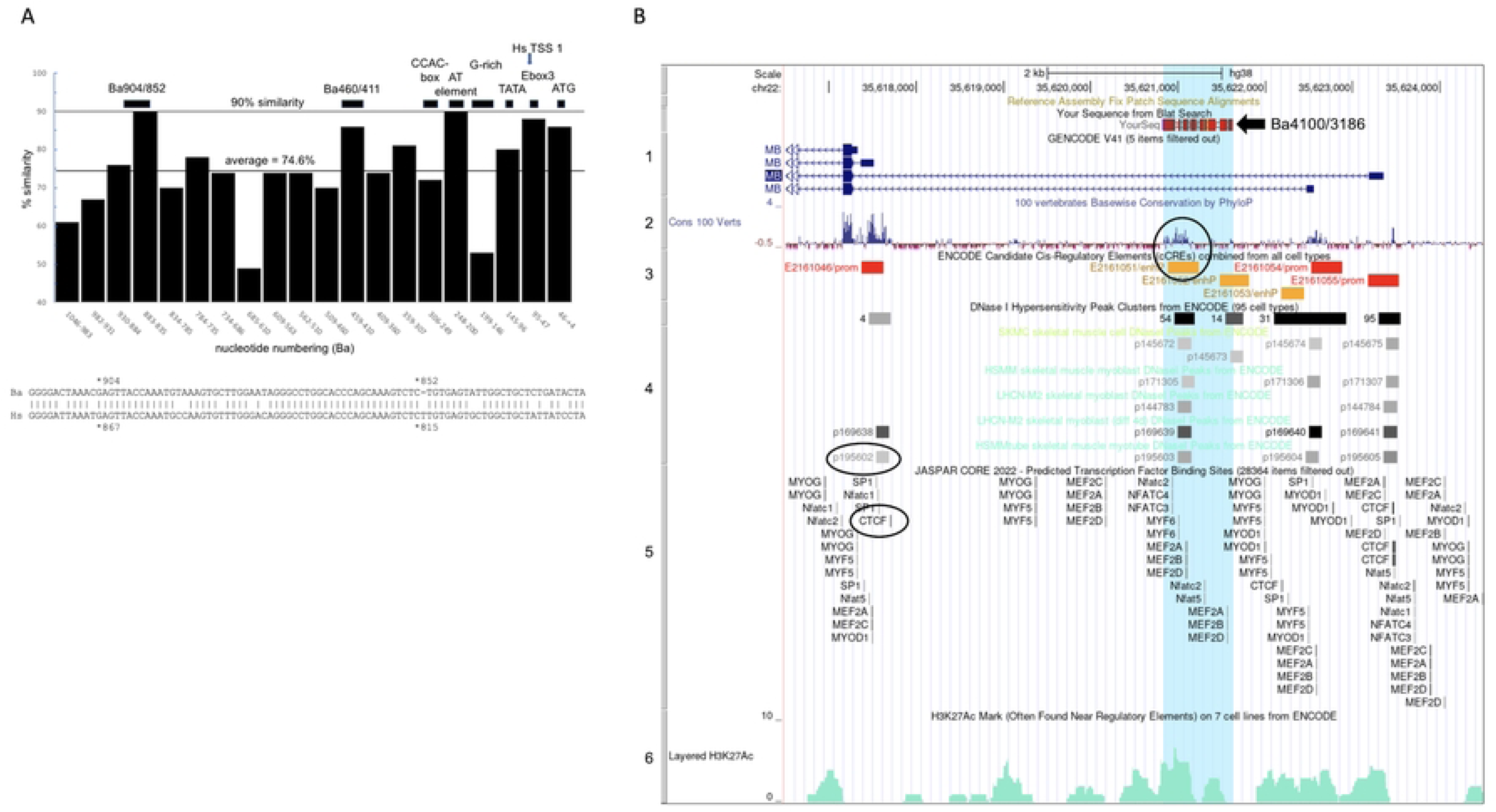
Deletion of the entire AT element is required to impact Ba promoter activity. (A) Alignment of AT element sequences from Hs and Ba, with predicted transcription factor binding sites conserved in both species (rVISTA) and expressed in muscle (S7 File) shown above. The two E-boxes and the core of the MEF2 binding site [31] are underlined. Below are the mutations tested. rVISTA predicts that Ba MEF mut eliminates MEF2 binding but does not impact binding of other factors; Ba E-box1 mut eliminates MYOD, E12, and MYOG binding but has no impact on MEF2 or E-box2 binding; Ba E-box2 mut eliminates DEC and TFE binding without affecting E-box1 or MEF2 binding. (B) Activity of AT element mutations. To determine which samples differ from control, ATswap was used for the comparisons; samples indicated by ** had p <0.01 (see S3D File for details). ATswap: AT swap, mean = 101% of control. MEFmut: Ba MEF mut, mean = 88% of control. ΔAT: Ba ΔAT, mean = 68% of control, p <0.01. Ebox1mut: Ba E-box1 mut, mean = 90% of control. Ebox2mut: Ba E-box2 mut, mean = 91% of control. Ebox3mut: Ba E-box3 mut, mean = 57% of control, p <0.01. (C) Alignment of the E-box3 region from Hs and Ba (E-box3 is addressed in the text below), with predicted transcription factor binding sites expressed in muscle (S7 File) shown above. Binding site sequences in lower case were identified independently on the Hs and Ba sequences by LASAGNA; binding sites in upper case were identified as conserved by rVISTA. MYOD binding is also predicted by MATCH. The human binding site sequences are shown for simplicity; the T → C difference at nt 59 does not prevent binding for any of the transcription factors shown (numbering from the ATG is the same in Hs and Ba for this region). Note that all three E-boxes received the same mutation: CAnnTG ◊ GAATTC.

In all, three nucleotides differ between Ba and Hs within the AT element (Fig 3A). When these three nucleotides were mutated in the Ba promoter to match the Hs sequence (AT swap) the resulting activity was unchanged relative to the intact Ba710 control (Fig 3B).

Previously, mutation of the AT-rich core of the MEF2 site in the Hs promoter (“MEF2 mut”) was shown to reduce expression to about 20% of wild-type [14]. To test the MEF2 site in the Ba promoter for function, we reproduced this mutation in the Ba promoter to create Ba MEF mut (Fig 3A). In Ba, this mutation reduced expression only slightly, to 88% of control (Fig 3B). Mutations in the two E-boxes (Ba E-box1 mut and Ba E-box2 mut, Fig 3A) also reduced activity only slightly, to 90% and 91% of control, respectively (Fig 3B). In contrast, when we deleted the entire AT element (Ba ΔAT, deletion (Δ) of Ba248/222 and replacement with a GAATTC sequence, Fig 3A), expression is reduced to 68% of control (Fig 3B). When multiple comparisons were made between AT swap, Ba MEF mut, Ba E-box1 mut, Ba E-box2 mut, and Ba ΔAT, the only mutation with a statistically significant difference from the others was Ba ΔAT (Fig 3B, S3D File). This indicates the MEF2 site does not play the dominant role in the Ba gene that it does in the Hs gene, and each of the three components of the Ba AT element ar edispensable, such that mutation of any individual component has a minimal effect, but deletion of all three reduces expression by more than 30%.

#### CCAC-box

The CCAC-box is a GC-rich region 5’ of the AT-element, first defined as a “myoglobin upstream regulatory element” (MbURE) critical for transcriptional activity [13]. The CCAC-box is reported to bind the ubiquitous transcriptional activator SP1 [16] but may also bind other transcription factors [14,17,32] (S4A File). In the Hs gene, the AT element and CCAC-box interact synergistically to drive high levels of muscle-specific expression [15,18].

Mutation of the CCAC-box in the Hs gene reduces activity to about 10% of wild-type [14]. However, the sequences around the CCAC-box are poorly conserved across species (S4A File). This raises the question of whether the Ba CCAC-box is a fully functional ortholog of the Hs CCAC-box. To test this, we reproduced the “CCAC mut 3” of Bassel-Duby et al. [14] in the Ba promoter, creating Ba CCAC mut (Fig 4A). We find that Ba CCACmut reduces activity to 85% of control (Fig 4B), not a statistically significant reduction (S4C File).

**Fig 4.**
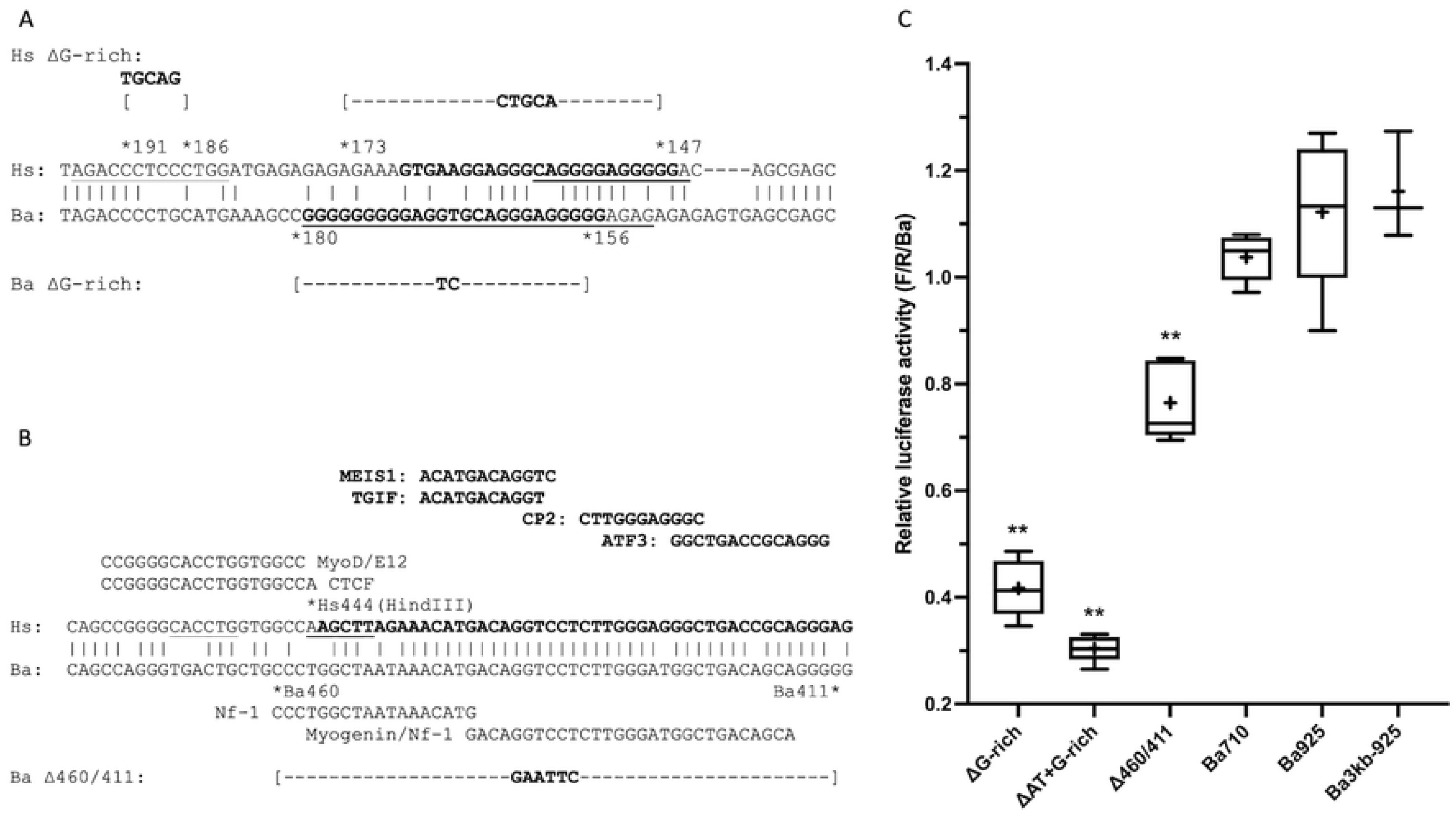
The Ba CCAC-box has little detectable activity in differentiated C2C12 cells. (A) Alignment of the CCAC-box region from Hs and Ba, with predicted transcription factor binding sites conserved in both species (rVISTA) and expressed in muscle (S7 File) shown above. An NFAT site not found by rVISTA but previously identified [84] is underlined. The 5’ SP1 site targeted by Ba ΔSP1-CCAC is in bold. The region targeted by the Ba CCAC swap is boxed in the Hs sequence. Below are the mutations tested. rVISTA predicts that Ba CCAC mut eliminates the PATZ1/MAZR and SP1 sites but a new nonconserved site is created (agctCCTCCCcgg in Ba); this new site is not created in CCAC mut 3 [14]. (B) Activity of CCAC-box mutations. For analysis of statistical significance in this figure, Ba ΔSP1-CCAC was used for comparison; samples indicated by ** had p <0.01 (see S4C File for details). CCAC mut: Ba CCAC mut, mean = 85% of control. ΔCCAC: Ba ΔCCAC, mean = 110% of control, p <0.01. ΔSP1-CCAC: Ba ΔSP1-CCAC, mean = 92% of control. ΔCCAC-AT: Ba ΔCCAC-AT, mean = 59% of control, p <0.01. CCACswap: CCAC swap, mean = 119% of control, p <0.01. CCAC+ATswap: CCAC+AT swap, mean = 171% of control, p <0.01. CCACswap vs.CCAC+AT swap: p <0.01. (C) Comparison of Ba410 to Ba410 ΔCCAC. 410: Ba410, mean = 75% of control. 410ΔCCAC: Ba410 ΔCCAC, mean = 82% of control.

We next made a 67 nt deletion removing the CCAC-box and a surrounding C-rich stretch containing multiple CAC motifs (Ba ΔCCAC, replacement of Ba317/251 with a GAATTC sequence; Fig 4A). Ba ΔCCAC was found to increase activity to 110% of control (Fig 4B) rather than decreasing it as expected. We noted a conserved predicted SP1 site (rVISTA) immediately 5’ of this deletion (Fig 4A) and hypothesized that the increased activity may be due to bringing this SP1 site closer to the AT element, and this may mask a possible loss of activity. However, extending the deletion to remove the SP1 site (Ba ΔSP1-CCAC, Ba334/251) still only slightly reduced expression, to 92% of control (Fig 4B).

In the Hs gene, the CCAC-box and AT element activate expression synergistically [15,18]. To test whether combining Ba ΔCCAC with Ba ΔAT would display a synergy that would reveal an effect of the Ba CCAC-box, we made Ba ΔCCAC-AT (contiguous deletion of Ba317/222, S1B File), which encompasses both regions. This larger deletion of 96 nt had a small but statistically significant reduction in activity compared to the Ba ΔAT deletion alone by two-tailed t-test (Ba ΔAT = 68% of control, Fig 3B; Ba ΔCCAC-AT = 59% of control, Fig 4B; Ba ΔAT (*M* = 0.677, *SD* = 0.062), Ba ΔCCAC-AT (*M* = 0.592, *SD* = 0.045); *t*(8) = 2.500, *p* = 0.037).

We also tested whether introducing the Hs CCAC-box and AT element sequences together into the Ba promoter would be sufficient to confer the high level of expression characteristic of the Hs promoter. We mutated the Ba CCAC-box to match the Hs sequence (CCAC swap, Fig 4A); these changes increased activity to 119% of control (Fig 4B). When the nucleotide changes of the CCAC swap were combined with the three nucleotide changes of the AT swap (CCAC+AT swap), the activity increased synergistically to 171% of control.

Much of the previous work on the Hs MB gene, including the construction of CCAC mut 3 [14], used a clone ending at a HindIII site at Hs444 (corresponding to Ba 457). We hypothesized that redundant or compensating sequences present between Ba457 and Ba710 may be masking the influence of the Ba CCAC box. We therefore tested the Ba ΔCCAC mutation in the context of a Ba promoter truncated to Ba410 (Ba410 ΔCCAC, deletion of Ba317/251, Fig 4C). The Ba410 endpoint leads to a reduction in expression to 75% of control; Ba410 ΔCCAC is expressed at 82% of control (S4D File), not a statistically significant difference by two-tailed t-test (Ba410 (*M* = 0.755, *SD* = 0.020), Ba410ΔCCAC (*M* = 0.824, *SD* = 0.142); *t*(8) = 1.076, *p* = 0.313).

Taken together, these data indicate that the Ba CCAC-box is not a major contributor to the activity of the Ba gene, as it is in the human gene. Deleting 84 nt from the CCAC-box region (Ba ΔSP1-CCAC) has little effect on expression. In contrast, introducing the human CCAC-box sequences into the Ba gene led to a statistically significant increase in expression, and the human CCAC-box interacts synergistically with the human AT element within the context of the Ba gene.

#### Human AT element and CCAC-box deletions

The above experiments prompted us to validate the ability of our assay system to detect deletions of the AT element and CCAC-box, so we made corresponding deletions in the Hs671 promoter. We find that deletion of the Hs AT element (Hs ΔAT, replacement of Hs246/219 with GAATTC, Fig 3A) reduced expression to 24% of the full length Hs671 promoter (Fig 5). This is similar in magnitude to the human MEF2 mut [14], but greater than the reduction to 68% seen with Ba ΔAT. Deletion of Hs CCAC (Hs ΔCCAC, replacement of Hs307/249 with GAATTC, Fig 4A) reduced expression to 65% of Hs671 (Fig 5). Hs ΔCCAC has a much more modest effect than the reduction to ~10% seen for the human CCAC mut 3 [14], but still greater than seen in any of our Ba CCAC-box mutations. Therefore, we conclude that the C2C12 assay system is capable of showing significant reductions in activity in response to the ΔAT and ΔCCAC deletions, and that the Ba CCAC-box and AT elements do not share the levels of activity previously shown in the Hs gene.

**Fig 5.**
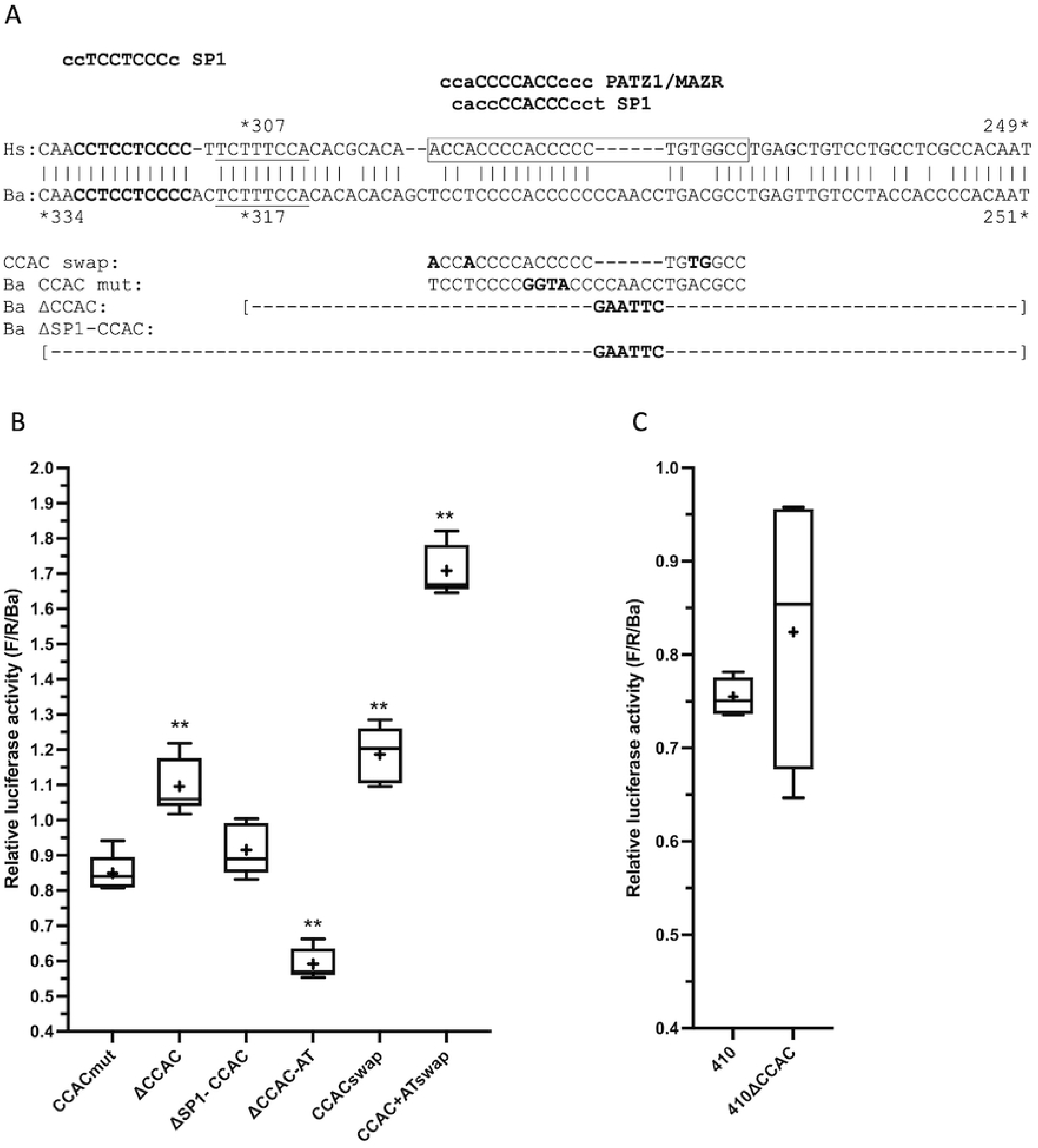
The human AT element and CCAC-box have detectable activity in differentiated C2C12 cells. Activity of deletions in Hs671, normalized to Ba710 (F/R/Ba). Samples that differ from Hs671 are indicated by asterisks (* p <0.05, ** p < 0.01; see S5B File for details). Hs671: mean = 12.7-fold Ba710. Hs ΔAT: mean = 3.1-fold Ba710 (24% of Hs671), probability of a difference from Hs671 (p) = <0.01. Hs ΔCCAC: mean = 8.2-fold Ba710 (65% of Hs671), p = 0.019. Hs ΔG-rich: mean = 10.0-fold Ba710 (79% of Hs671), probability of a difference from Hs671 is not significant.

#### E-box3

In addition to the two E-box sequences in the AT element, a third conserved E-box exists, in the 5’UTR of MB exon 1 (Fig 1). This E-box, termed E-box3 [28], is predicted (rVISTA, LASAGNA) to bind several transcription factors expressed in muscle (Fig 3C). E- box3 was explored in mice [28] and found to repress expression in a muscle-specific fashion, that is, mutation increased expression. In contrast, we find that the same mutation used for E-box1 and E-box2 decreased activity to 57% of control (E-box3 mut, Fig 3B, C). Therefore, Ba E-box3 is an activating element, in contrast to its’ action as a repressive element in the mouse gene.

### Two novel regulatory regions are revealed in studies of the Ba gene

We used bioinformatics to guide further analysis of the Ba MB gene, and uncovered two additional functional regions, a G-rich sequence at Ba179/155, and a conserved sequence at Ba449/412.

#### G-rich sequence

A search conducted for transcription factor binding sites (rVISTA) revealed that SP1 binding sites occur in only two places in the Ba gene: in the vicinity of the CCAC-box (Fig 4A) and in a strikingly G-rich sequence 3’ of the AT element (80% G over 25 nucleotides, Ba179/155, Fig 6A). The G-rich sequence is flanked at its’ 3’ side by a repeated GAGA motif [33]. We therefore targeted this region for further study. A deletion of 25 nucleotides in the G-rich sequence (Ba ΔG-rich, Fig 6A) strongly reduced expression to 42% of control (ΔG-rich, Fig 6C). Deletion of the GAGA motif (BaΔGAGA, deletion of Ba155/145) had no effect (S6A File).

**Fig 6.**
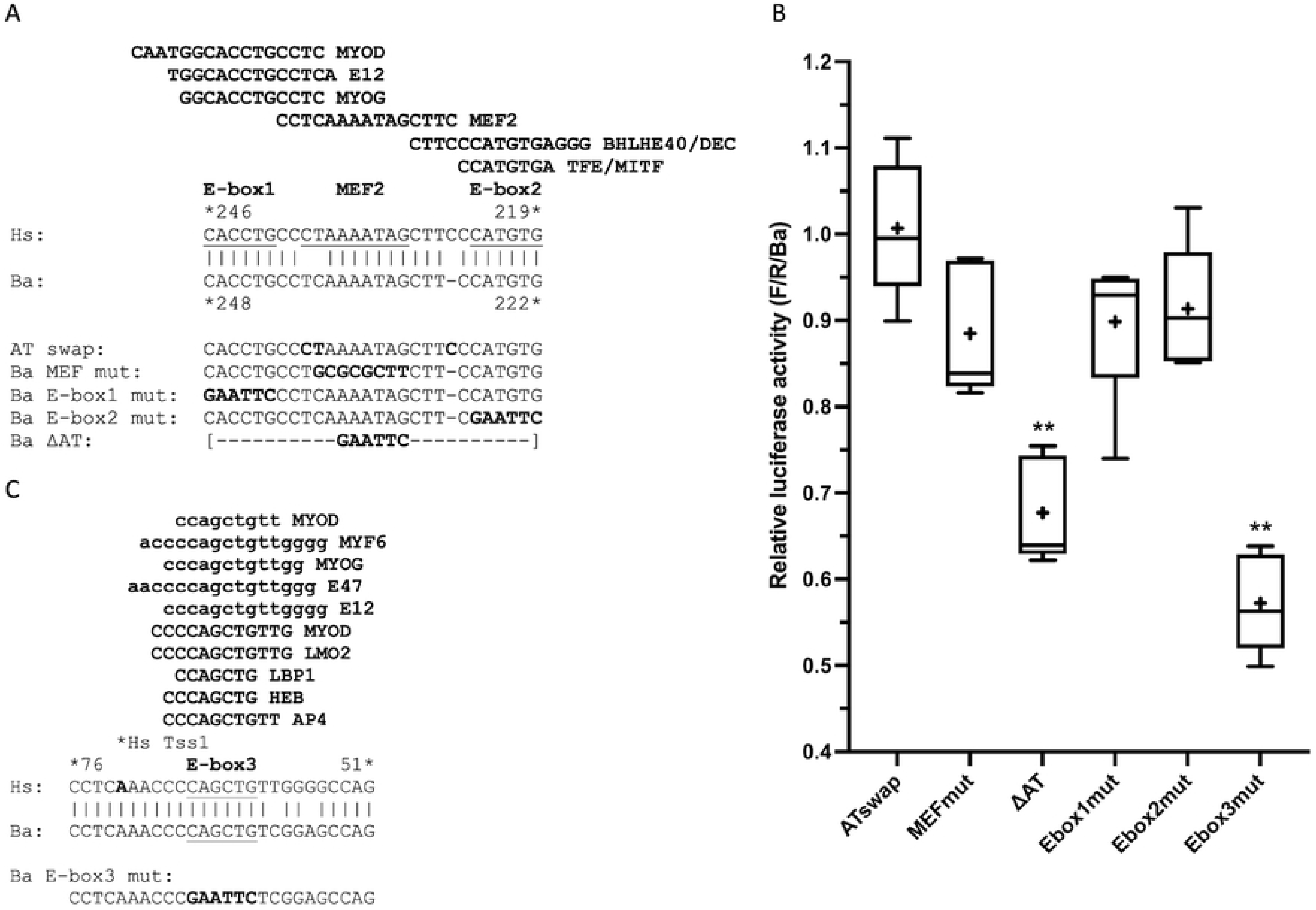
Sequences and activities of novel regulatory sequences in the Ba gene. (A) Alignment of the region around the Hs and Ba G-rich sequences. The G-rich sequences Hs168/146 and Ba179/155 are in bold font. The sequences predicted (rVISTA) to bind SP1 are underlined; none of these predicted binding sites is conserved between Hs and Ba. An additional predicted SP1 binding site (MATCH) 5’ of the Hs G-rich sequence (Hs195/183) is also underlined. The Ba ΔG-rich mutant replaces Ba180/156 with a TC sequence; the Hs ΔG-rich mutant replaces Hs191/186 with a TGCAG sequence and Hs173/147 with a CTGCA. Analysis of both mutant sequences (rVISTA, MATCH) predicts no SP1 binding sites. (B) Alignment of the Ba449/412 conservation between Hs and Ba. In the Hs sequence, the 5’ end of the DNaseI hypersensitive site p195602 is in bold text and a nonconserved flanking CTCF binding site is indicated (from UCSC Genome Browser [20], see Fig 7B, Tracks 4 and 5). Above, conserved (rVISTA) sites for transcription factors expressed in muscle (S7 File) are indicated in bold. An E-box (CACCTG) is underlined and nonconserved MYOD and E12 (rVISTA) sites are indicated. A HindIII site (AAGCTT, at position −373 of Devlin et al. [13]) is shown for reference. In the Ba sequence, nonconserved NF1 and composite MYOG/NF1 sites (LASAGNA) are indicated. The Ba Δ460/411 deletion replaces Ba460/411 with a GAATTC sequence. (C) Activity of mutations in the novel Ba regulatory regions. For analysis of statistical significance in this figure, Ba710 was used for comparison; samples indicated by ** had p <0.01 (see S6C File for details). ΔG-rich: Ba ΔG-rich, mean = 42% of control, p <0.01. ΔAT+G-rich: Ba ΔAT + ΔG-rich, mean = 30% of control, p <0.01. Δ460/411: Ba Δ460/411, mean = 76% of control, p < 0.01. 710: Ba710, mean = 104% of control. Ba925: Ba925, mean = 112% of control. 3kb-925: Ba3kb-925, mean = 116% of control.

In the Hs promoter, the CCAC-box is proposed to bind SP1, which then interacts synergistically with transcription factors bound to the AT element [16,18]. The relative inactivity of the Ba CCAC-box led us to hypothesize that this G-rich cluster of predicted SP1 sites may be the functional substitute for the Hs CCAC-box. We therefore combined the AT element deletion, Ba ΔAT, with the G-rich sequence deletion, Ba ΔG-rich, to give Ba ΔAT+ΔG-rich (Ba Δ248/222+Ba Δ180/156, S1B File). The combined deletions reduced expression even further, to 30% of control (Fig 6C). The difference between Ba ΔG-rich and Ba ΔAT+ΔG-rich is statistically significant by two-tail t-test (Ba ΔG-rich (*M* = 0.417, *SD* = 0.054), Ba ΔAT+G-rich (*M* = 0.304, *SD* = 0.025); *t*(8) = 4.255, *p* = 0.003). Since Ba ΔAT alone reduced expression to 68%, and Ba ΔG-rich reduced expression to 42%, it appears that the AT element and G-rich sequence act additively (68% of 42% = 28%), but synergistic interaction is not observed.

The Hs gene also shows predicted (rVISTA) SP1 binding sites in only two places: the CCAC-box and a similarly positioned G-rich sequence (70% G over 23 nt, Hs168/146, Fig 6A). A repeated GAGA motif is present, but in the Hs gene it is 5’ of the G-rich sequence. Deletion of the Hs G-rich sequence and an adjacent SP1 site (Hs ΔG-rich, Fig 6A) does not lead to a statistically significant decrease in expression (Fig 5 and S5B File), although there is a trend toward lower expression (S5A File). Therefore, this region may have weak function, but is clearly not a major determinant of activity in the Hs promoter as it is in the Ba promoter.

#### Conserved sequence at Ba449/412

Deletion of the Ba promoter to Ba410 reduced expression to 75% of Ba710 (Fig 4C), indicating the region between Ba410 and Ba710 houses additional regulatory elements. In a survey of sequence similarity between Ba and Hs across ~1,000 nt 5’ of the MB coding region, an above average region of similarity occurs between Ba460/411 (Fig 7A). Within this region, a stretch of 36/38 conserved nucleotides occurs at Ba449/412 (95% identity, Fig 6B). Since high conservation of non-coding sequence between species can reflect regulatory function [34–37], we explored this sequence further.

**Fig 7.**
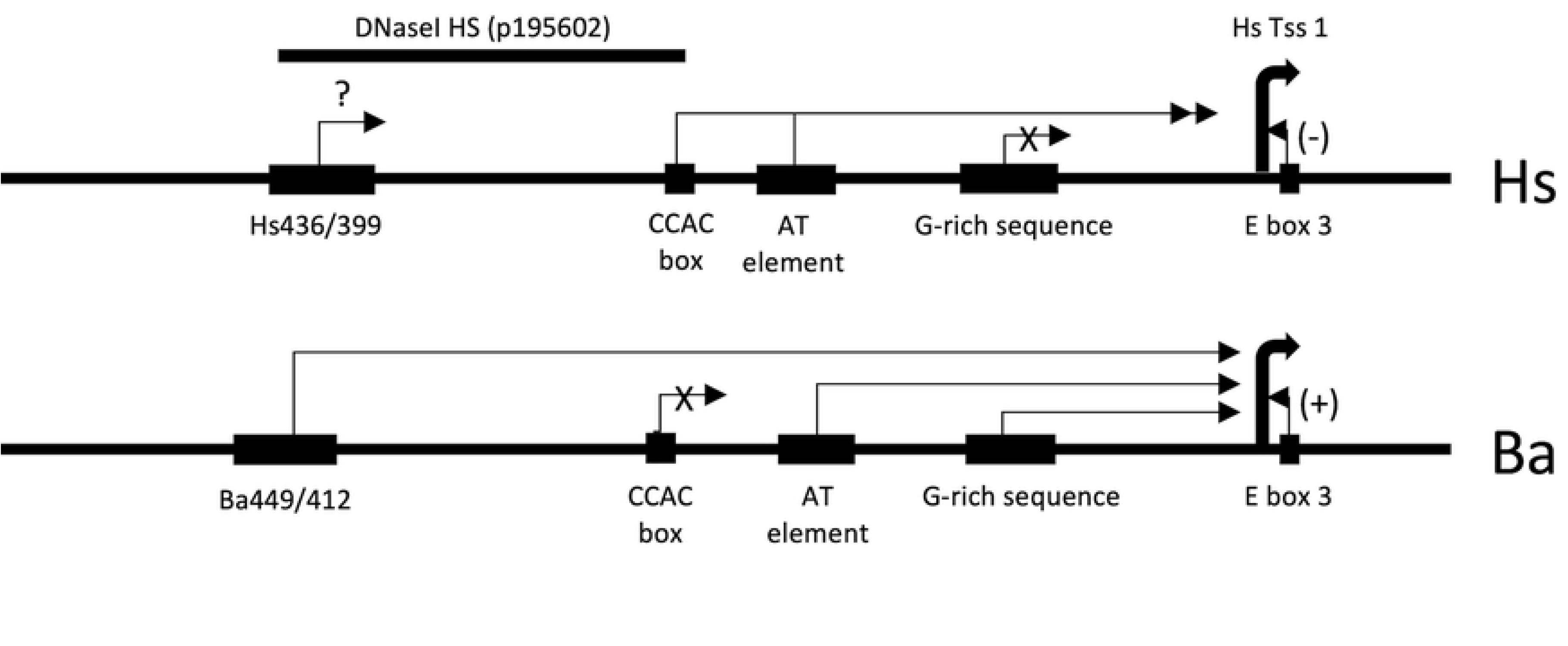
Two distal regions of high sequence conservation have no discernible activity. (A) Graph of percent sequence similarity between Hs and Ba from nucleotide +4 (A of ATG = +1) to Ba1046 and Hs996, in blocks of 50 nts relative to the Hs sequence. The average nucleotide similarity across this region is 74.6%. Shown above are regulatory landmarks described in this work. Below is a BLASTn alignment of 100 nt of Ba and Hs sequence around the Ba904/852 conserved region. (B) Presentation of 8,000 nt of human chromosome 22 from the UCSC Genome Browser [20,21] showing the 5’ end of the human myoglobin gene and about 7,000 nt of the 5’ flanking region. Selected browser tracks are numbered at left. Above Track 1 the Ba4100/3186 region tested as the putative “Ba3kb” enhancer is indicated. The circle spanning Tracks 2 and 3 encloses a sequence similarity peak and the computationally predicted Candidate Cis-Regulatory Element “e2161051/enhP”. Circled in Tracks 4 and 5 are the DNaseI hypersensitive site (“p195602”) and the CTCF site shown in Fig 6B. Track 1: the 4 most-proximal of the nine known transcription start sites; the one at the top is TSS 1, the major transcription start site, at coordinate chr22:35,617,329; transcription is right to left. Track 2: Sequence conservation across 100 vertebrates. Track 3: Computationally predicted candidate cis-regulatory elements based on ENCODE data. Track 4: DnaseI hypersensitive sites in five muscle cell line samples. Track 5: Selected transcription factor binding sites predicted by JASPAR. Track 6: Histone H3 lysine27 acetylation. This UCSC Genome Browser view can be accessed at: https://genome.ucsc.edu/s/csackerson/chr22%3A35%2C616%2C500%2D35%2C624%2C499

In the Hs gene, this region has several notable features (Fig 6B): First, this region is the 5’ end of a DNase I hypersensitive region [38] in muscle cells. Second, consistent with this region being a boundary between active (DnaseI hypersensitive) and inactive chromatin, it is flanked by a predicted CTCF [39] binding site (JASPAR, LASAGNA). Third, a cluster of binding sites for transcription factors expressed in muscle is predicted (rVISTA, S7 File), and this cluster is conserved in the Ba promoter. Fourth, this region has an additional E-box identical in sequence to E-box1 (CACCTG) that is predicted (rVISTA) to bind MYOD and E12. In addition, in a comparison between human and mouse sequences, successful alignment between the two species ends near the HindIII site at Hs444 (S6D File). Matches are not found further 5’, consistent with this region being the 5’ end of the MB regulatory region.

The DNaseI hypersensitive status of the Ba gene is unknown, and Ba is not predicted to bind CTCF (LASAGNA), nor does it have the E-box. However, it is predicted to bind nuclear factor 1 (NF1) at two adjacent sites within this region (rVISTA, LASAGNA). NF1 is expressed in muscle (S7 File) and, like CTCF, contributes to boundary functions in chromatin [40]. We made a deletion of this sequence in the context of Ba710 (Ba Δ460/411, Fig 6B), which reduced expression to 76% of control (Δ460/411 in Fig 6C), essentially identical to the reduction in activity seen with the truncated Ba410 promoter (Fig 4C). Deletion of this region in the Hs gene was not conducted.

### Distal conserved sequences do not increase activity

One possible explanation for the low activity of the Ba gene compared to the Hs gene is that activating regulatory elements exist that are not included in the Ba710 construct. Using DNA sequence conservation as a guide, we identified a region at Ba904/852 and Hs867/815 with 91% identity (48/53 nt) between the species (Fig 7A). The only other region with similarly high conservation is at the AT element (59/64 matches, 92% identity). We extended the cloning to a Ba925 endpoint to include this conserved region in our analysis. Although the Ba925 construct increased mean activity to 112% of the Ba710 control (Fig 6C), this is not a statistically significant increase (S6C File).

To gain a genome level view of conservation across the MB locus, we queried the UCSC Genome Browser [20] for information from the ENCODE project [41]. The ENCODE Project has catalogued various markers of candidate regulatory elements in the human genome such as DNA sequence conservation, transcription factor binding, DNase I hypersensitivity, and histone modifications. Our search is shown in Fig 7B. A region of high DNA sequence conservation between multiple species is located ~3.6 kb 5’ of the major transcriptional start site in the human sequence. This coincides with a “candidate Cis-Regulatory Element” (cCRE) designated “E2161051/enhP” (Fig 7B, Tracks 2 and 3). This region is DNase I sensitive in the five muscle cell lines tested (Fig 7B, Track 4), and shows moderate histone H3K27 acetylation modifications (Fig 7B, Track 6). cCRE E2161051 also contains predicted binding sites for MYF6, MEF2 and NFAT transcription factors (Fig 7B, Track 5).

cCRE E2161051 maps by alignment to Ba3699/3347. Examination of the surrounding region showed that sequence conservation between Ba and Hs remains high (~80%) for approximately 300 nt further 5’. We cloned a ~900 nt fragment that includes the putative cCRE and the additional conserved region (Ba4100/3186, “Ba 3kb”), and inserted it 5’ of the Ba925 vector to create Ba3kb-925. In three paired transfections (Fig 6C, S6B File), Ba3kb-925 and Ba925 were essentially identical in activity by two-tailed t-test (Ba925 (*M* = 1.122, *SD* = 0.141), Ba93kb-925 (*M* = 1.161, *SD* = 0.101); *t*(6) = 1.409, *p* = 0.696). Therefore, despite strong bioinformatics clues of activity of this region, no activity was observed in our assay system.

## DISCUSSION

### Transcription driven by Cetacean 5’ flanking regions is only about 6% that of humans

Cetaceans are notable for the high levels of myoglobin in their skeletal muscle. Traditionally, transcriptional levels have been considered a dominant (although not sole) determinant of protein levels [17,42,43]. We therefore hypothesized that Cetacean myoglobin genes would be highly active transcriptionally, and that study of their transcriptional mechanisms may provide insights into myoglobin transcriptional control, and regulatory evolution.

However, we found that, on average, the MB genes from Cetacean species were only 6% as active, on average, as the human gene. We expanded our survey to the closely related, terrestrial Artiodactylan species. Here we included elk to have a non-domesticated Artiodactylan species, since myoglobin levels in domesticated species may be driven by breeding in response to market forces [44]. Again, observed activities were similar to those seen for whales, and much lower than that of humans. Moving further out phylogenetically, we tested horses and found that the horse gene is >5-fold more active than the human gene, and >90-fold more active than the Cetaceans and Artiodactylans. Last, we tested a carnivore, dogs, and obtained an activity ~2-fold higher than that of the Cetacean and Artiodactylan mean (Table 1, Fig 2A, S2 File).

This survey demonstrated that transcriptional levels from myoglobin genes can vary greatly, and do not correlate well with myoglobin protein levels. For example, myoglobin protein in human type 1 skeletal muscle fibers is ~4.5 mg/g [45], whereas minke whales contain ~20 mg/g [12]. Yet, despite the 4-5-fold higher protein level in the whale tissue, the minke whale gene is only 8% as active in our assays. There are several possible explanations for this observation.

One possibility is that our cloning missed critical positive cis-acting elements. However, expanding the size of the regulatory region tested to Ba925 did not increase activity to a statistically significant degree (Fig 6C). Further, the inclusion of the candidate cis regulatory element “E2161051/enhP” also did not increase expression levels (Fig 6C and Fig 7). In the work of Devlin et al. [13] on the human gene, ~2000 nt of 5’ sequence was tested for activity. Reducing the size of this tested region did not cause a decrease in activity until they encountered what they termed the myoglobin upstream regulatory element (MbURE), beginning at Hs331 (our numbering; −261 in [13]). Therefore, although distal regulatory elements may exist, available data does not support their presence.

A second possibility is that physiological signaling, for example from muscle contraction [22,23], hypoxia [23,46], nitric oxide [47], or a lipid-rich diet [48,49], might induce increased expression. Our experiments were conducted in differentiated, cultured C2C12 cells without the manipulation of calcium-induced pathways, hypoxia, or other physiological pathways, so we cannot address these possibilities. However, we do observe high activity of the human and horse genes in our assay system, so we can at least conclude that constitutive expression driven by developmental differentiation mechanisms is low from the Cetacean and Artiodactylan genes.

A third possibility is that post-transcriptional mechanisms work to increase protein levels in the absence of increased transcriptional levels. Such mechanisms might include increased mRNA stability, high translational levels, and stability of existing protein. A recent genomic survey of the correlation between mRNA and protein levels in skeletal muscle finds that transcription levels and protein levels do not correlate for myoglobin [24], so post-transcriptional regulation is indicated. This observation is also consistent with work on emperor penguins, where it was found that myoglobin protein levels increased to a greater degree than mRNA levels in adults versus chicks [50]. A high level of protein stability has evolved in myoglobins from diving mammals, so protein stability may be a major contributor to the high tissue levels found [25,26].

The high myoglobin level found in diving mammals creates a challenge in that myoglobin can aggregate at high concentrations [51], and such aggregates may lead to the destruction of the myoglobin [52]. This problem may be solved in aquatic mammals through the evolution of high net electrostatic surface charge [53] which inhibits aggregation [51]. One might speculate that high protein stability, and further protein evolution to inhibit aggregation may have coevolved with low transcriptional activity. In contrast, horse MB is 4-fold less stable than minke whale MB [25], and aggregates at lower concentrations than sperm whale myoglobin [54]. Thus, the higher gene expression levels seen in horses may conversely have evolved to compensate for a higher rate of protein turnover.

### The minke whale gene has substituted a G-rich sequence for the CCAC-box

We have dissected the 5’ flanking regulatory region of the minke whale (Ba) MB gene, guided by previous studies in humans and mice, and clues from bioinformatics. We find numerous differences between the minke gene and the human gene (Fig 1). The most striking is the relative inactivity of the minke whale CCAC-box ortholog. Instead, we propose that a G-rich sequence serves a similar function, cooperating with the AT element to activate transcription.

Considerable evidence supports the importance of the CCAC-box in the human gene. Devlin et al. [13] found that deletion of 5’ sequences to Hs331 (−261 in [13]) had little effect on expression levels, but further deletion to Hs275 (−205 in [13]) reduced expression to 12%. This deletion defined the MbURE, which contains the CCAC-box. Internal deletion of the region between Hs444 (HindIII, −373 of [13]) and Hs275 in the context of ~1,000 nt of 5’ sequence completely abolished expression. In the work of Bassel-Duby et al. [14], a subtle mutation of Hs285/282 from ACCC to GGTA (CCAC mut 3) in the context of the Hs444 (HindIII) 5’ sequence reduced expression to less than 10% of the wild-type sequence. Further, Bassel-Duby et al. [15] and Grayson et al. [18] demonstrate synergistic interaction of the CCAC-box and the AT element. Our own results support these observations. Changing the Ba CCAC-box to match the human sequence increased expression, and combining the CCACswap with the ATswap increased expression synergistically (CCACswap and CCAC+ATswap, Fig 4B), even though the ATswap by itself had no effect on expression (ATswap, Fig 3B).

In contrast, we find that our Ba ΔSP1-CCAC (Ba334/251) has no effect on expression from the Ba gene. This deletion corresponds to Hs323/249, very similar to the 5’ deletion of Devlin et al. [13] from Hs331to Hs275 that defined the MbURE. We also recreated the CCAC mut 3 mutation in the Ba gene, with no effect. The only evidence we have for significant activity of the Ba CCAC-box is when its’ deletion is combined with deletion of the AT element. The combination, ΔCCAC-AT (Fig 4B) shows a slight reduction in activity when compared to ΔAT alone (Fig 3B), but this deletion is quite large, removing 96 nt.

The human CCAC-box is reported to bind the ubiquitous activating transcription factor, SP1 [16]. We noted predicted binding of SP1 not only to the C-rich (G-rich on the other strand) CCAC-box 5’ of the AT element, but also to a G-rich (C-rich on the other strand) sequence 3’ of the AT element in both the human and minke whale genes. When we tested the Ba G-rich sequence for function, we found that its’ deletion (ΔG-rich) strongly reduced expression (Fig 6C). In contrast, the human G-rich sequence is nonessential (Fig 5). Combining the deletion of the Ba G-rich sequence with deletion of the Ba AT element (ΔAT+G-rich, Fig 6C) reveals an additive effect between the two. We therefore propose that, in the minke whale gene, the G-rich sequence has become substituted for the CCAC-box, but the synergy seen in the Hs gene between the CCAC-box and the AT element has been lost.

Both the CCAC-box and the G-rich sequence are poorly conserved between humans and minke whales (Fig 4A, 6A, 7A). Repetitive sequences are prone to rapid change, through DNA slippage during replication. [55–58]. The presence of these runs of G-C nucleotides may have facilitated the functional swap between the CCAC-box and the G-rich sequence through their evolutionary flexibility. It is well established that cell-type specific gene expression is controlled by combinations of transcription factors bound at promoters and enhancers [59]. Both the human and minke genes are predicted to bind MYOD, MYOG, E12 and MEF2 factors at the AT element (Fig 3A), and SP1 at their GC-rich sequences (Fig 6, S4A File), so one might expect similar activity from the two enhancers if they function simply through the accumulation of critical transcription factors. However, the order, distance, and orientation of the transcription factors bound to an enhancer, what has been termed “enhancer grammar”, can matter in some cases [60]. Thus, despite a similar constellation of factors bound in the Ba and Hs genes, changing the order in which these transcription factors bind may be responsible for the lower transcriptional output, perhaps through the loss of synergy as mentioned above.

We have presumed above that the CCAC-box and the G-rich sequence function primarily by binding SP1. SP1 binding to the CCAC-box of humans and minke whale is robustly predicted (S4A File). However, experimental evidence of SP1 binding to the human CCAC-box is not direct. Grayson et al. [16] describe a reporter plasmid containing a sequence from the human muscle creatine kinase gene similar to the MB CCAC-box, which responds synergistically to cotransfected SP1- and MEF2A-expressing vectors in *Drosophila* SL2 cells. Further, also in the human muscle creatine kinase gene, SP1 is shown to be present in a complex with MEF2 bound to an AT element-like sequence. Although direct binding assays between the human myoglobin CCAC-box and SP1 were reported, the data is not shown.

Alternative explanations exist for the activity of the human CCAC-box and minke G-rich sequence. Runs of G have been shown to form G-quadruplex structures that are highly enriched at promoters [61] where they may mediate a variety of functions including epigenetic regulation. G-quadruplex structures have also been shown to bind MYOD [62]. Indeed, MYOD binding to a non-E-box site in the human CCAC-box is predicted (S4A File); MYOD binding to the minke whale CCAC-box is not predicted. In addition, this predicted MYOD binding is not found when the CCAC mut 3 mutation of Bassel-Duby et al. [14] is examined (rVISTA). Last, the human CCAC-box binds a nuclear factor of 40 kD [14], the approximate size of MYOD (see, for example, [63]; 34.5 kD in Uniprot [64]); in contrast, SP1 has a molecular weight of approximately 80.7 kD [64,65]. Thus, the human CCAC-box may have functions beyond the binding of SP1 that contribute to its’ relative importance, compared to the minke CCAC-box. Similarly, the minke G-rich sequence may function through mechanisms beyond SP1 binding.

### Functioning of the MEF2 and E-box sites have evolved between the human and minke whale genes

At the core of the AT element is a MEF2 binding site [14,15,66] flanked on both sides by E-boxes (Fig 3A). An additional E-box occurs immediately 3’ of the transcriptional start site (Fig 3C). The functions of these sites differ between humans and minke whales.

In previous studies of the human AT element [14,15] mutation of the MEF2 site reduced expression to 20% of wild-type. We used the same nucleotide changes in the Ba gene, but expression was reduced only to 88% of control, a reduction that was not statistically significant. This mutation is predicted to no longer bind MEF2, and this is the only predicted MEF2 binding site in the Ba710 sequence.

This difference is unlikely to be due to a lack of MEF2 activity in our assay system. Previous studies used the mouse skeletal muscle cell line Sol8 [14] or mouse and rat heart muscle [15], while we used C2C12 cells. However, MEF2 has been shown to be expressed in C2C12 cells by three days of differentiation [67]. Also, electrophoretic mobility shift assays comparing DNA binding activities of nuclear proteins from sol8 and C2C12 myotubes appear identical [18]. Last, although MEF2 is activated by increases in intracellular calcium via the calcineurin phosphatase [68], MEF2 can activate gene expression in C2C12 cells after differentiation in the absence of specific manipulation of calcium flux [69,70]. Therefore, MEF2 is likely to be both present and capable of activating gene expression in our assay system.

Mutations of the minke whale E-boxes also had different results from those reported for the human E-boxes. In the human gene, E-box1 was previously shown to be nonessential, in keeping with our results, but E-box2 was shown [14] to be required for full activity. Mutation of the minke whale E-box2 had no effect in our experiments. Only when the entire AT element is deleted is there an effect on expression (Fig 3B). It is possible that the components of the minke whale AT element serve redundant functions such that loss of any one component has no effect.

The minke whale E-box3 may also play a role in these differences. Mutation of the minke E-box3 showed a significant decrease in expression (Fig 3B), whereas E-box3 mutation in the human promoter is reported [28] to increase expression. E-box3 is of the “symmetrical” type (CAGCTG, [71]) capable of binding MYOD as a homodimer, as well as MYOD-E2A heterodimers [72]. Binding to MYOD is robustly predicted for E-box3 from both humans and minke whales (Fig 3). Interactions between MYOD bound to E-box3 and MEF2 [73–75] may mask the loss of E-box1 or 2, or E-box3 may interact with E-box1 or 2 to activate expression in the absence of MEF2 [76]. Intriguingly, MYOD-E12 heterodimers bound to an E-box can interact with SP1 and serum response factor (SRF) to activate gene expression [77, reviewed in 78]. As described, SP1 binding is broadly predicted in the minke promoter-proximal sequences. Binding of SRF is predicted (LASAGNA, MATCH, rVISTA) in only two places in the Ba710 minke whale sequence: within E-box2 (CCATGTGAGG, Ba228-219) and immediately upstream of E-box3 (CCCTTTAGGGCCA, Ba94-82). SRF is not predicted (rVISTA) to bind the Hs671 human gene. Thus, the mechanisms of activation of the minke whale gene may differ in numerous ways from the human gene.

### Novel regulatory sequences are found in the minke gene

Our explorations of minke whale MB regulation revealed two functional regions that had not been noted in studies of the human gene. The first is the G-rich sequence discussed above. The second is a conserved sequence at Ba449/412 (tested as a larger deletion of Ba460/411). The features of this region in the human gene include the 5’ end of a DNase I hypersensitive region and a CTCF binding site (Fig 7B). These observations are consistent with this region being the 5’ boundary of a functional regulatory element [79,80]. CTCF is not predicted to bind the Ba sequence, but two NF1 binding sites are predicted (Fig 6). NF1 has been shown to have insulator activity, delineating a boundary between chromatin domains [40], much like CTCF. Thus, although the minke region is not predicted to bind CTCF, and the human region is not predicted to bind NF1, the functional consequences may be the same. The Ba460/411 deletion removes not only the NF1 sites, but also binding sites for muscle transcription factors conserved between humans and minke whales, and caused a significant loss of activity. It is unknown why the activity of this region was not uncovered in previous studies of the human gene [13]. Further exploration of the role of this region in MB gene control may be productive.

### Limitations of the study

The approaches taken in these studies have several inherent limitations. First, the assays for function were conducted by transient transfection into a cultured cell line growing in vitro. One limitation of this approach is that the transfected gene lacks its’ native genomic context such as neighboring genes, transcriptionally associated domains, and regulation by chromatin [81]. However, transfection has been a mainstay of gene expression studies for nearly 50 years [82]. As we have demonstrated, transfection can provide valuable first steps in understanding previously unknown aspects of gene regulation.

A second limitation is that cultured cells growing in vitro similarly lack context, in this case the complex physiology of intact muscle tissue. For example, two previously identified NFAT transcription factor binding sites [17,83,84] are found in the human promoter. The NFAT transcription factor couples muscle activity to the expression of proteins involved in contractility [85,86], and activates myoglobin expression [84]. One of these binding sites is conserved in the minke whale promoter, immediately 5’ of the CCAC-box; this site was deleted in the Ba ΔSP1-CCAC mutant without any effect (Fig 4B). The other site at Hs764 (−690 of Chin et al. [84]) is not conserved, but two others are predicted (LASAGNA) at Ba588 and Ba810. The Ba925 construct contains both sites; deletion to Ba511 removes them both and still results in expression at 106% of control (S6B File). Since changes in calcium concentration that would accompany muscle contraction in the intact tissue is not recapitulated in our assays, the potential effect of the NFAT transcription factor is not addressed. Similarly, other physiological influences would not be detected.

A third limitation of our studies is our reliance on bioinformatics-based signals of function, such as transcription factor binding predictions and inter-species DNA sequence conservation. Despite the promise of such approaches, the majority of “hits” must necessarily be false positives [reviewed in 87]. To minimize this false positive problem, we have given the most credibility to transcription factor binding predictions that are conserved between humans and minke whales, an evolutionary distance in excess of 50 million years. Nonetheless, they are still “predictions” and without direct experimental validation they are best considered hypotheses. Similarly, inter-species DNA sequence conservation increases in predictive value when multiple species are aligned (see Supporting Information for several examples). The conserved region at Ba460/411 was found to have function, but the conservation at Ba904/852 and at the −3kb region did not correlate with an effect on transcription in our assays (Fig 6C, Fig 7).

### Conclusions

Our most striking observation in these studies is the low constitutive expression of the minke whale myoglobin gene, in contrast to the high myoglobin protein levels seen in adult muscle tissue. We propose that the low transcription is the result of multiple evolved regulatory differences between the human and minke genes. These regulatory differences may have coevolved with an increased stability of the myoglobin protein [53,54,88], leading to high myoglobin protein levels despite low constitutive transcriptional activity. These findings also point to the importance of the induction of transcription by physiological influences such as exercise [84,86] and diet [89] in the ontogeny of the high muscle myoglobin protein levels seen in Cetaceans.

## Materials and Methods

### Species survey tissue and genomic DNA samples

Total genomic DNA was isolated from tissue samples using DNeasy Blood & Tissue Kit (Qiagen, catalog # 69504).

Cetacean tissue samples were collected from stranded, deceased animals by member organizations within the National Marine Mammal Stranding Network, The Hawaii Pacific University Stranding Program (HPUSP), the Oregon Marine Mammal Stranding Network (OMMSN), the International Fund for Animal Welfare (IFAW), and the Alaska Stranding Network (ASN) under the authority of a NMFS Stranding Agreement issued to each of the cooperating organizations by the Office of Protected Resources, National Oceanic and Atmospheric Administration. Laboratory use of the tissues was conducted under a Marine Mammal Parts Handling Authorization, provided by regional offices of National Marine Fisheries Service. As this prior review had been conducted by experts in this field further ethical review by the cooperating institution, California State University Channel Islands, was not required.

Human DNA was isolated from a saliva sample provided by CS. Cow DNA was isolated from locally purchase beef. Elk DNA was isolated from elk steak purchased from Basspro.com (catalog # 1568645).

Horse (catalog # GE-170) and dog (catalog # GD-150M) DNA was purchased from Zyagen.com.

### Myoglobin gene cloning and preparation of plasmids used for transfection

See S8 File for details of clonings and primers used for each reported construct. In general, MB gene sequences were isolated through two rounds of PCR using Accuprime Pfx DNA polymerase (Invitrogen, cat # 12344-024). The first round used as the 5’ (forward) primer “bosF1”, a degenerate primer designed from an alignment of human, horse, and cow sequences (beginning at a nucleotide equivalent to Ba1024 for reference). The 3’ (reverse) primer “stenR3” was designed from the dolphin *Stenella attenuata* exon 1 sequences (beginning at the equivalent of nt Ba+65). The PCR products of this first round were typically complex, so a second round of PCR was carried out using nested primers. In the second round, various 5’ primers were used (S8 File). Two 3’ primers, “stenR1” and “stenR2”, were used depending on which gave a unique product; these primers were designed from *S. attenuata* exon 1 sequences (beginning at Ba+3 or Ba+51, respectively). The blunt-ended round two product was sufficiently pure for cloning into pIBI31 cut with SmaI and selected as white colonies on Xgal plates. The clone was moved into pGL4.10[luc2] (Promega, catalog # E6651) with XhoI and NcoI. The result of this strategy is the inclusion of 30 nt of pIBI31 polylinker sequence at the 5’ end of the pGL4.10 clone. All the species tested have their ATG in a CCATGG sequence recognized by NcoI, so the 3’ end was a direct fusion of the cloned ATG translational initiation sequence to the firefly luciferase ATG of pGL4.10.

As genomic sequences became publicly available, subsequent clonings and manipulations used 5’ forward primers with an XhoI site at its 5’ end to allow direct cloning into pGL4.10 without extraneous polylinker sequences at their 5’ end, as detailed in S8 File. This difference was not seen to affect expression in a systematic way.

The pSV-Rluc renilla luciferase internal standard vector was derived from psiCHECK-2 (Promega, catalog # C8021) by digestion with EagI to remove the firefly luciferase gene, leaving the renilla luciferase gene driven by the SV40 promoter.

All pGL4.10 derivatives used for transfections were verified by sequencing.

For transfection, plasmids were purified on QIAprep Spin Miniprep columns (Qiagen, catalog # 27104) with an extra wash with PB binding buffer, and quantitated on a NanoDrop ND-1000 spectrophotometer.

### Cell culture, transfection, and luciferase assays

Mouse C2C12 cells [90] used were obtained from ATCC (catalog # CRL-1772), except three experiments for which the cells used were obtained from the laboratory of Barbara Wold at CalTech (originally from ATCC), as indicated in S1 Table.

For routine passage, cells were plated at 4×10^5^ cells in 60mm tissue culture plates in “growth medium”: DMEM (high glucose, without pyruvate; Thermo-Fisher cat # 11965092) with 10% fetal bovine serum (Gibco cat # A3160501); no antibiotics were used. Cells were passaged every two days. All experiments were performed with cultures that had gone through fewer than 10 passages since receipt.

For transfection, cells were plated in 12-well tissue culture plates at 1×10^5^ cells per well in 1.2 ml growth medium. Immediately after plating, the transfection mix was prepared and added to the cells, within about 1 hour of plating. Transfections used 1 μg pGL4.10-derived reporter plasmid plus 1 ng pSV-Rluc renilla luciferase for normalization (see below) in a total volume of 50 μl with DMEM (no serum). 5 μl Polyfect Transfection Reagent (Qiagen, catalog # 301105) was added, vortexed for 10 seconds, incubated 10 minutes, and diluted with 300 μl growth medium; the entire volume was added to each well. In practice, duplicate wells were transfected from a master mix scaled up 2.2-fold from the above quantities. After 24 hours, the media was replaced with DMEM with 2% horse serum (Fisher, catalog # 301105) and ITS (Gibco, catalog # 4140045) at 1:1000 dilution (“differentiation medium”); this is day zero of differentiation. Media was refreshed every day until the cells were harvested.

In some initial experiments included in the species surveys, each sample was transfected in triplicate, and differentiation was carried out for 6 days. These parameters were changed to 4 days of differentiation to avoid cell detachment, and samples were transfected in duplicate for all later experiments. Differentiation was extensive after 4 days of differentiation judged by morphology and expression of the slow isoform of the myosin heavy chain (S9 File). These changes were not seen to affect the results after normalization to a Ba710 control included in duplicate in each transfection plate (that is, each 12-well culture plate included 2 wells of the Ba710 control plus 5 experimental samples in duplicate). In all cases replications were averaged, and this average represents an n=1.

Assays used the Dual-Luciferase Reporter Assay System (Promega, catalog # E1910). Each transfected well was lysed into 200μl Passive Lysis Buffer, and 50μl was used to measure firefly activity with 50 μl LARII, followed directly by 50 μl Stop&Glo to measure renilla luciferase activity, on a FilterMaxF5 microplate reader (Molecular Devices) using a 96-well opaque white microplate.

### Data analysis and statistics

Luciferase activity was processed as follows: firefly luciferase counts were first normalized to the co-transfected renilla luciferase control to give “F/R”. F/R was also calculated for two Ba710 samples included in each plate as a control and averaged. Then, for each transfected plate, F/R for each experimental sample was divided by the average of the Ba710 control to give “F/R/Ba”; descriptions of activity as “x% of control” refer to the F/R/Ba value. For Fig 6D, independent experimental sets of Ba710 transfections were processed in the same way to generate an independent experimental Ba710 sample.

Statistical analysis and graphing was done with GraphPad Prism (version 9), using standard parametric tests. For multiple comparisons, ANOVA was followed by Tukey’s HSD test [91]. Data presented in each figure was analyzed as a set, using one of the samples with a mean ~1.0 for post-hoc Tukey comparisons, as specified in the figure legend. In cases where pairwise comparisons were made, a two-tailed t-test was used; p < 0.05 was considered significant, with further consideration for multiple identical tests (alpha - k/n).

### Bioinformatics

We searched for transcription factor binding sites in Ba sequences using the online search programs: MATCH [92] using the “muscle_specific.prf” profile, LASAGNA [93], and rVISTA [94]. rVISTA queries two sequences to look for conserved binding sites. Transcription factor predictions in the human sequence presented in the UCSC Genome Browser interface used JASPAR [95]. Expression of transcription factors in muscle cells described in figure legends and S7 File is according to https://www.genecards.org [96] and www.proteinatlas.org [97]. Dot-plot alignment (S6D File) is from PipMaker [98].

## Acknowledgements

We are indebted to the National Marine Mammal Stranding Network, The Hawaii Pacific University Stranding Program (HPUSP), the Oregon Marine Mammal Stranding Network (OMMSN) the International Fund for Animal Welfare (IFAW) and the Alaska Stranding Network (ASN) for supplying the Cetacean tissues. We are also indebted to CSUCI and the Biology Department, and especially its’ Chairs, for generous provision of space and support; and the Biology Support staff, especially Mike Mahoney, Cathy Hutchinson, and Jessica Dalton. Last, we recognize the many students that have contributed to this project over the years in the context of the Biology Department Independent Research (Biol 494) course.

## Supporting Information Captions

**S1 File. Supplement to Fig 1.**

**A** Flow chart of the experiments. Created with BioRender.com. See S9 File for photographs of differentiated C2C12 cells after 4 days.

**B** Schematics of the internal deletions described in the text. To the right is the name of the deletion and in parentheses the nucleotides deleted in each case. Except for the two Ba410 derivatives, deletions are in the context of Ba710.

**S2 File. Full data set for Table 1.**

Data is expressed as F/R/Ba (see Materials and Methods). Four species not shown for simplicity in Table 1 or Fig 2 are included here: *Balaenoptera musculus*(Bm706), *Megaptera novaeangliae* (Mn704), *Orca orcinus* (Oo703), and *Sus scrofa* (Ss657). Data indicated by a superscript “x” is from clones inserted directly into the XhoI and NcoI sites of pGL4.10 without extraneous polylinker sequences. Data generated in C2C12 cells from the Wold lab are indicated by a superscript “w”; all other data was generated in C2C12 cells purchased directly from ATCC. The average of the seven Cetacean species shown is 0.801; including the three Artiodactylan species yields an average of 0.782 and this number is used for Fig 2B (CetArt). “SEM” is standard error of the mean.

**S3 File. Supporting information for Fig 3.**

**A** Alignment of the AT element (E-box1--Mef-2--E-box2): Hash marks indicate identity with the human (Hs) sequence. Most Cetaceans (Mn, Er, Bm, Dc, Oo) and the Artiodactylans (Bt, Cc) are identical to Ba through the AT element; Pp has a variant E-box1 (CACATG instead of CACCTG).

**B** Alignment of E-box3 region: Most species have the canonical CAGCTG sequence, except for the Odontoceti, Oo (*Orca orcinus*), Dc (*Delphinus capensis*), Tt (*Tursiops truncatus*), and Pp (*Phocoena phocoena*). The bottlenose dolphin, *T. truncatus*, is included because no published sequence is available for the common dolphin, *D. capensis*. Mm: *Mus musculus*. The variant sequence in the Odontoceti is not predicted by rVISTA to bind any of the transcription factors shown in Fig 3C.

**C** Full data set, average of duplicate wells, normalized as F/R/Ba. Equal variances confirmed, based on homogeneity of variances test (indicated here and below with an asterisk on the construct name). Analysis of variance (ANOVA) confirms a statistical difference between the samples (*F*(5,24) = 24.955, *p* <0.001). ANOVA was followed by the post-hoc Tukey HSD test [35].

**D** Tukey test data for Fig 3B

**S4 File. Supporting information for Fig 4.**

**A** Multiple species alignment of the CCAC-box region: The 10 nt CCAC-box [15] encompasses Hs289/280 and Ba297/288. Hash marks indicate identity with the human (Hs) sequence. Sp1 binding (bold type) to the Ba sequence is robustly predicted by rVISTA, LASAGNA, and MATCH, and to the Hs sequence by rVISTA, MATCH and JASPAR; Sp1 binding is also predicted by rVISTA for Pp, Ss, and Cf, but not for Bt or Ec. Other transcription factors that are predicted by rVISTA (but not necessarily conserved) and are expressed in muscle are shown: USF (underline), MYOD (wavy underline in the Hs sequence), AP2 (asterisks), and PPARalpha (double dashed underline).

**B** F Full data set, average of duplicate wells, normalized as F/R/Ba. Equal variances confirmed, based on homogeneity of variances test. ANOVA confirms a statistical difference between the samples (*F*(7,35) = 99.449, *p* <0.001). ANOVA was followed by the post-hoc Tukey HSD test.

**C** Tukey test for Fig 4B.

**D** Full data set for Ba410 and Ba410ΔCCAC, average of duplicate wells, normalized as F/R/Ba.

**S5 File. Supporting information for Fig 5.**

**A** Full data set, average of duplicate wells, normalized as F/R/Ba. Equal variances confirmed, based on homogeneity of variances test. ANOVA confirms a statistical difference between the samples (*F*(3,15) = 16.109, *p* <0.001). ANOVA was followed by the post-hoc Tukey HSD test. The data shown is derived from five transfected plates, allowing direct comparison of the constructs; note that although the data for HsΔG-rich is not statistically different from that for Hs671, in each case the HsΔG-rich samples are lower than the Hs671 values.

The same Hs671 data shown here was included in the data set in Table 1.

**B** Tukey test for Fig 5.

**S6 File. Supporting information for Fig 6.**

**A** Full data set, average of duplicate wells, normalized as F/R/Ba. Equal variances confirmed, based on homogeneity of variances test. ANOVA confirms a statistical difference between the samples (*F*(3,15) = 151.582, *p* <0.001). ANOVA was followed by the post-hoc Tukey HSD test.

**B** Full data set, average of duplicate wells, normalized as F/R/Ba. The data in heavy boxes is derived from three transfected plates, allowing direct comparison of the activities of Ba710, Ba925, and Ba3kb-925. Equal variances confirmed, based on homogeneity of variances test. Analysis by ANOVA fails to find a statistical difference between the samples (*F*(3,15) = 1.152, *p* =0.360). ANOVA was followed by the post-hoc Tukey HSD test.

**C** Tukey test for Fig 6C.

**D** Left: Dot plot alignment (PipMaker [98]) of 1000 nt of mouse (*Mus musculus*, Mm) MB sequence 5’ of the ATG (Y-axis) against 1000 nt of human MB sequence 5 of the ATG (X-axis). The diagonal dashed line plots similarities between the two sequences using default settings. The 5’ end of the similarities occurs at nts Hs474 and Mm422. The HindIII site at Hs444 is shown for reference.

Right: Dot plot alignment of 1000 nt of Ba MB sequence 5’ of the ATG (Y-axis) against 1000 nt of Hs MB sequence 5’ of the ATG (X-axis) shown for comparison.

**E** Multiple species alignment of conserved sequences at Ba460/411. Vertical hash marks indicate identity with the Hs sequence. The HindIII site (AAGCTT) is bold for reference. Transcription factor sites conserved (rVISTA) with humans and expressed in muscle are indicated: ATF3 (underlined), MEIS1-TGIF (double underline), CP2 (asterisk), NFE2L1/TCF11 (dashed line), TTF1 (dotted)

**S7 File. Sources for transcription factors’ expression in muscle.**

**S8 File. Supporting information for Materials and Methods.**

**A** Details of the clonings used for the species scan.

**B** Published sequence accession numbers.

**C** Primers used for clonings.

**D** Primers and polymerase reagents used for mutations and deletions.

**S9 File. Differentiation of C2C12 cells under transfection conditions.**

Differentiation of C2C12 cells over 4 days in differentiation medium. Top row: Phase contrast pictures of transfected cells. Bottom row: Untransfected cells stained for myosin heavy chain; primary antibody was mouse monoclonal anti-myosin (skeletal, slow) from Sigma-Aldrich (cat # M8421) at 1:1000 dilution; detection with horse radish peroxidase used Vector ImmPress kit (cat # MP-7402).

